# Development and validation of an SDA-500 *Anopheles stephensi* cell line for molecular studies

**DOI:** 10.64898/2026.08.14.744812

**Authors:** Siji Kavil, Dylan Jinmi, Luke Alphey, Michelle A. E. Anderson

**Affiliations:** Department of Biology, University of York, York, UK; York Biomedical Research Institute, Department of Biology, University of York, UK

**Keywords:** *Anopheles stephensi*, Mosquito cell line, Transfection, CRISPR-Cas9, Malaria

## Abstract

**Background:** Malaria control is increasingly challenged by the urban-adapted vector *Anopheles stephensi*, yet molecular and cellular tools for this species remain scarce, restricting functional genomic studies and the development of genetic control strategies. To help address this gap, we established a new embryo-derived *Anopheles stephensi* cell line.

**Results:** We generated and characterised a novel embryo-derived *Anopheles stephensi* (SDA-500) cell line capable of sustained growth *in vitro*. Species identity was confirmed by mitochondrial COI barcoding, and karyotypic analysis revealed a diploid chromosome complement with the presence of a Y chromosome, confirming that at least some cells are of male origin. Transfection conditions were optimized, with TransIT-PRO showing higher efficiency than Lipofectamine-based reagents. Using a dual-luciferase reporter assay, of several promoters tested the *Anopheles gambiae* polyubiquitin promoter exhibited the strongest and most consistent transcriptional activity in SDA-500 cells.

**Conclusions:** The SDA-500 cell line provides a stable and genetically validated *in vitro* platform that supports efficient transgene expression. This resource provides a useful system for functional genomics and molecular manipulation in *Anopheles stephensi* and is expected to facilitate studies of mosquito biology and contribute to the development of novel malaria control strategies.

## Background

Malaria is the most significant vector-borne diseases globally, in terms of human mortality. It is caused by protozoan parasites of the genus *Plasmodium* and transmitted through the bites of infected female *Anopheles* mosquitoes. Despite decades of global control efforts, malaria continues to impose a substantial burden on human health, particularly in sub-Saharan Africa (1). In 2023, the World Health Organization estimated approximately 263 million malaria cases and nearly 600,000 deaths worldwide, with over 90% occurring in this region (1,2). Young children are especially vulnerable due to limited acquired immunity and persistent disparities in access to diagnosis and treatment.

Among malaria vectors, *Anopheles stephensi* has gained increasing attention because of its ability to thrive in urban environments. Historically the primary vector of urban malaria in the Indian subcontinent and parts of the Middle East, this species has recently expanded its range into Africa. Since its first detection in Djibouti City in 2012, *An. stephensi* has spread across the Horn of Africa and into parts of West Africa, where its establishment has coincided with substantial increases in malaria cases (3,4). Its adaptation to artificial water containers and densely populated urban settings raises growing concern about the future of malaria transmission in rapidly expanding African cities (5).

Addressing the challenges posed by emerging malaria vectors such as *An. stephensi* requires improved molecular tools and experimental platforms. Advances in genetic technologies, including CRISPR-Cas9-based genome editing and gene-drive systems, offer promising approaches for vector control (6). The development and optimization of these technologies, however, depend on reliable *in vitro* systems that enable controlled investigation of gene expression, genome manipulation, and potentially mosquito–pathogen interactions (7,8).

Mosquito-derived cell lines provide practical and experimentally tractable platforms for such studies, allowing preliminary testing of genetic constructs and regulatory elements before in vivo experimentation (9,10). Although insect cell biotechnology has advanced considerably over the past several decades, well-characterized *Anopheles* cell lines remain limited with most existing lines derived from species such as *Anopheles gambiae* and *Anopheles albimanus* (11,12). Comparable resources for *An. stephensi* are scarce, particularly in the United Kingdom limiting regionally accessible platforms for functional genomic studies in this increasingly important vector species. The establishment of insect cell lines often relies on embryonic tissues, which contain developmentally plastic cells capable of differentiating into diverse cell types and forming stable cultures (9). However, the value of a newly established cell line ultimately depends on its suitability for downstream molecular applications, including transfection and genome manipulation (8,10). A key step in this characterization is the evaluation of promoter elements capable of driving reliable gene expression within the cells (13,14).

Here we report the development and initial characterization of a newly derived *An. stephensi* cell line from embryonic tissue from the SDA-500 strain. We evaluated the transfectability of this cell line using three commercially available reagents and examined the activity of promoter fragments commonly used in mosquito vector research – including the synthetic 3xP3 promoter (15) and *polyubiquitin* promoters derived from *Anopheles gambiae, Aedes aegypti*, and *Aedes albopictus* – using a dual-luciferase reporter system (16,17). Karyotype analysis was performed to assess chromosomal integrity, and the presence of the Y chromosome in at least some cells, was confirmed by PCR using As*GUY1*-specific primers (18). Together, these analyses provide an initial validation of this cell line as a platform for genetic and functional genomic studies in *An. stephensi*.

## Results

### Establishment and validation of the SDA-500 cell line

Primary cultures were initiated from surface-sterilized *An. stephensi* embryos and maintained in L-15 medium supplemented with 20% FBS (Foetal bovine serum), 10% TPB (Tryptose Phosphate Broth), 1% Penicillin-streptomycin and 1% Gentamycin. After approximately six months of weekly media replacement the culture vessels became confluent and were passaged. Subsequent passages exhibited a more stable morphology and consistent proliferation, indicating adaptation to in vitro conditions. After approximately 25 passages, the cells showed consistent growth and appearance, requiring passaging twice a week at a 1:5 ratio, confirming the successful establishment of an immortalised cell line. The established cell line exhibited predominantly epithelial-like morphology (Fig. 1) and remained viable during routine passages.

**Figure 1.**
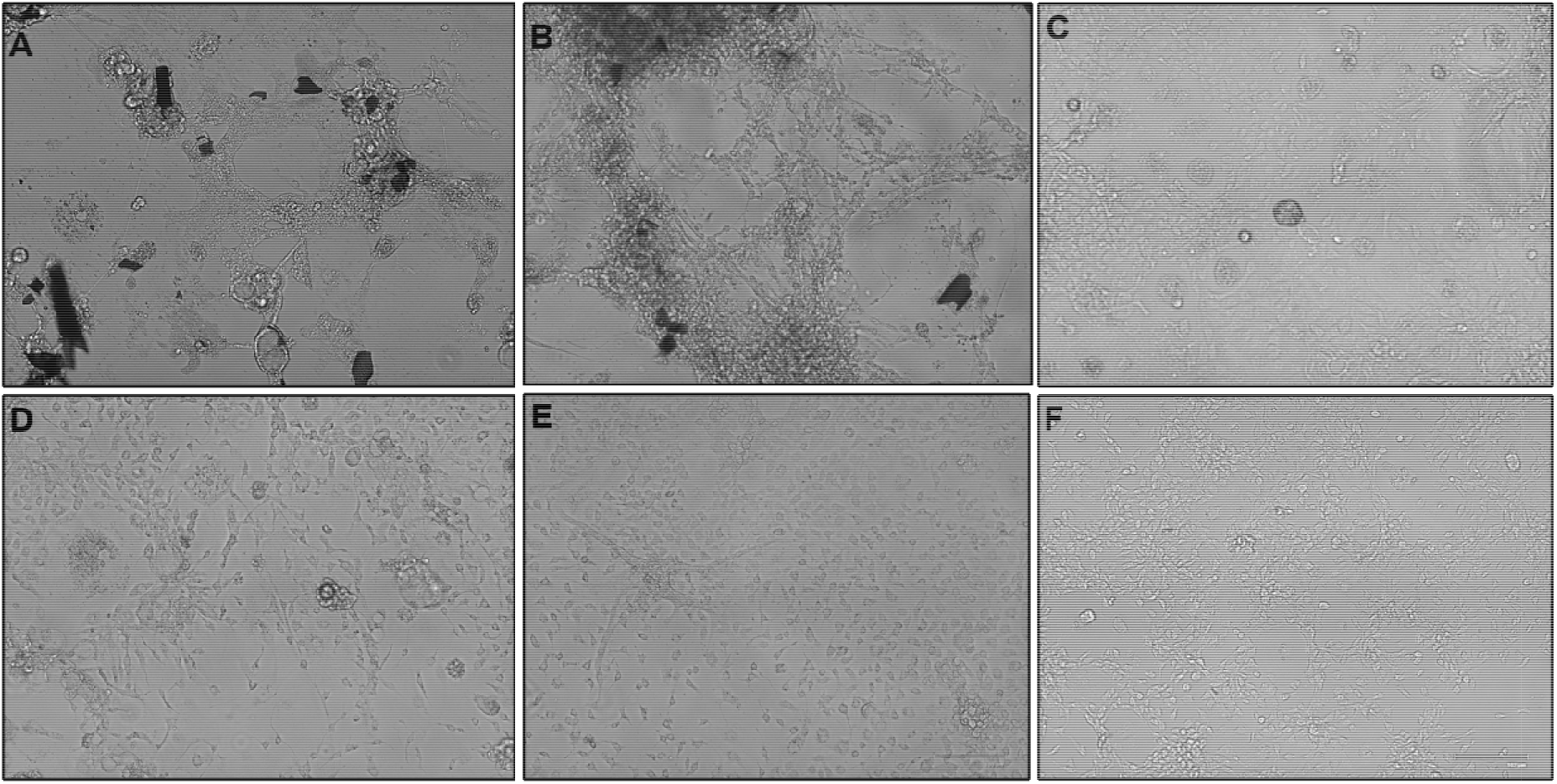
Sequential establishment and stabilization of the *Anopheles stephensi* primary cell line under *in vitro* culture conditions. Brightfield micrographs showing progressive establishment and adaptation of the *An. stephensi* primary cell culture. Cells maintained in T25 flasks exhibited gradual attachment and proliferation after 2 weeks (A), 3 months (B), and 6 months (C) of culture. The first subculture passage was performed upon attainment of confluency approximately 6 months after culture initiation. Images from passages P2 (D), P15 (E), and P27 (F) demonstrate progressive morphological stabilization and sustained proliferative growth of the established cell line. Brightfield images were acquired using a Floid Cell Imaging Station (Life Technologies) at a magnification of 20X.

### Species confirmation of the *An. stephensi* cell line by COI barcoding

To verify the species identity of the *Anopheles stephensi* cell line, a 720 bp fragment of the mitochondrial cytochrome c oxidase subunit I (COI) gene was amplified using the standard barcoding primers LCO1490 and HCO2198 and subjected to Sanger sequencing (19). Representative amplification and sequence confirmation are shown in **Additional file 1: Figure S1**. The resulting sequences were quality filtered, assembled, and queried against the NCBI nucleotide database using BLASTn. The top-scoring hit corresponded to *Anopheles stephensi*, exhibiting 99.55% sequence identity across the full length of the amplicon, with 98% query coverage and an E-value of 0.0. No other species in the database showed comparable similarity, supporting accurate species identification and excluding potential misidentification with other *Anopheles* species.

### Karyotyping and molecular confirmation of the Y chromosome in *An. stephensi* cells

Metaphase chromosome spreads revealed a consistent diploid chromosome number of 2n = 6 in cultured *An. stephensi* cells (Fig. 2A-C), in agreement with previously reported karyotypes for this species (20,21). Among the six chromosomes, one was consistently smaller and morphologically distinct, indicative of a Y chromosome (Fig. 2B). These observations were reproducible across multiple metaphase spreads.

**Figure 2.**
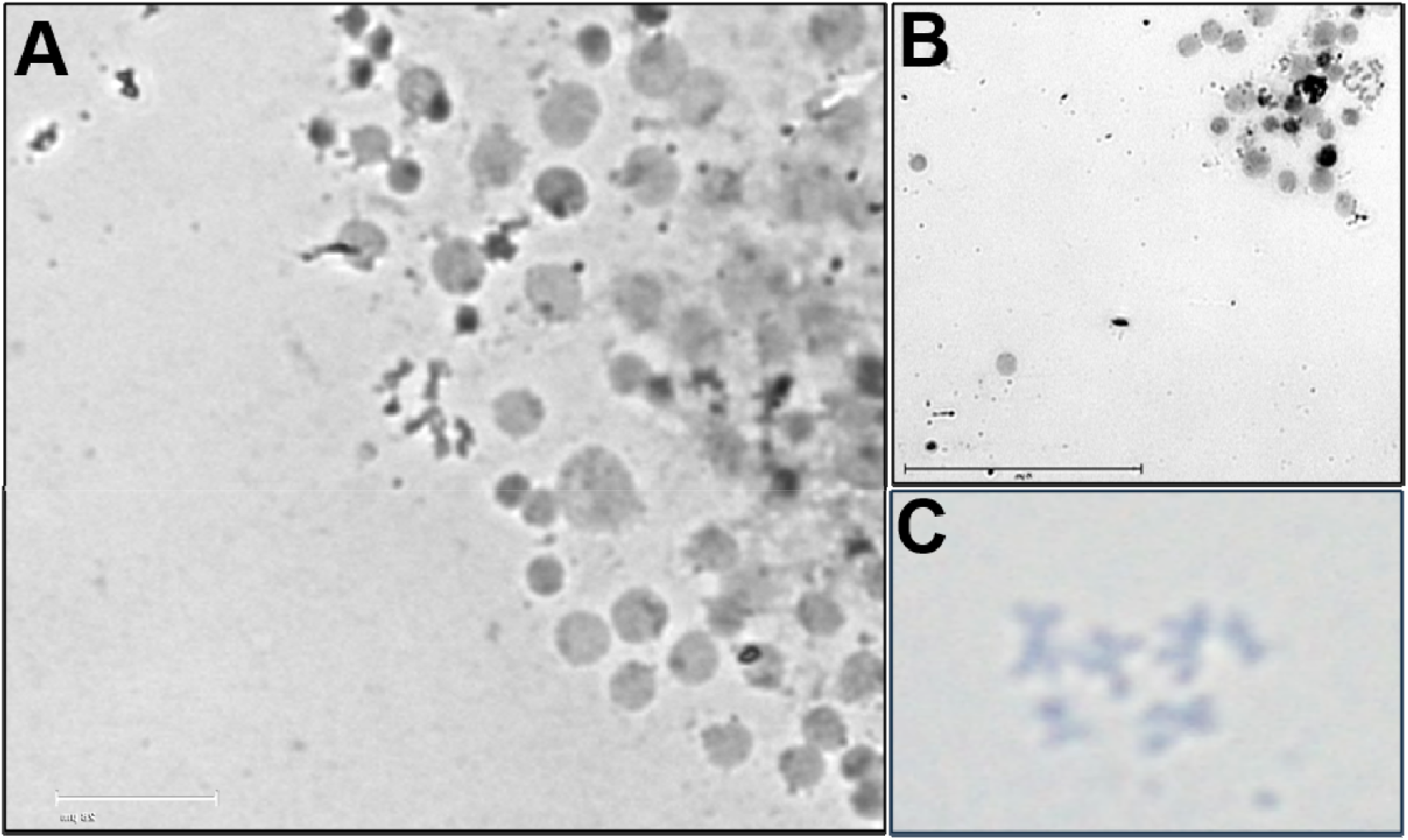
Karyotype of *An. stephensi* cells. Raw metaphase spread (A), Sister chromatid separation during mitotic metaphase (B). Images taken with EVOS M7000 imaging system at a magnification of 40x. Condensed metaphase chromosomes (C); Screenshot from Zeiss Observer 7 microscope at 40X magnification.

To verify the identity of the putative Y chromosome, PCR amplification was performed using *AsGUY1*-specific primers, which target a Y-linked gene known to be exclusively present in male *An. stephensi*. Amplification yielded a distinct PCR product of the expected size (Fig. 3), confirming the presence of the Y chromosome in at least some cells of the cultured cell line.

**Figure 3.**
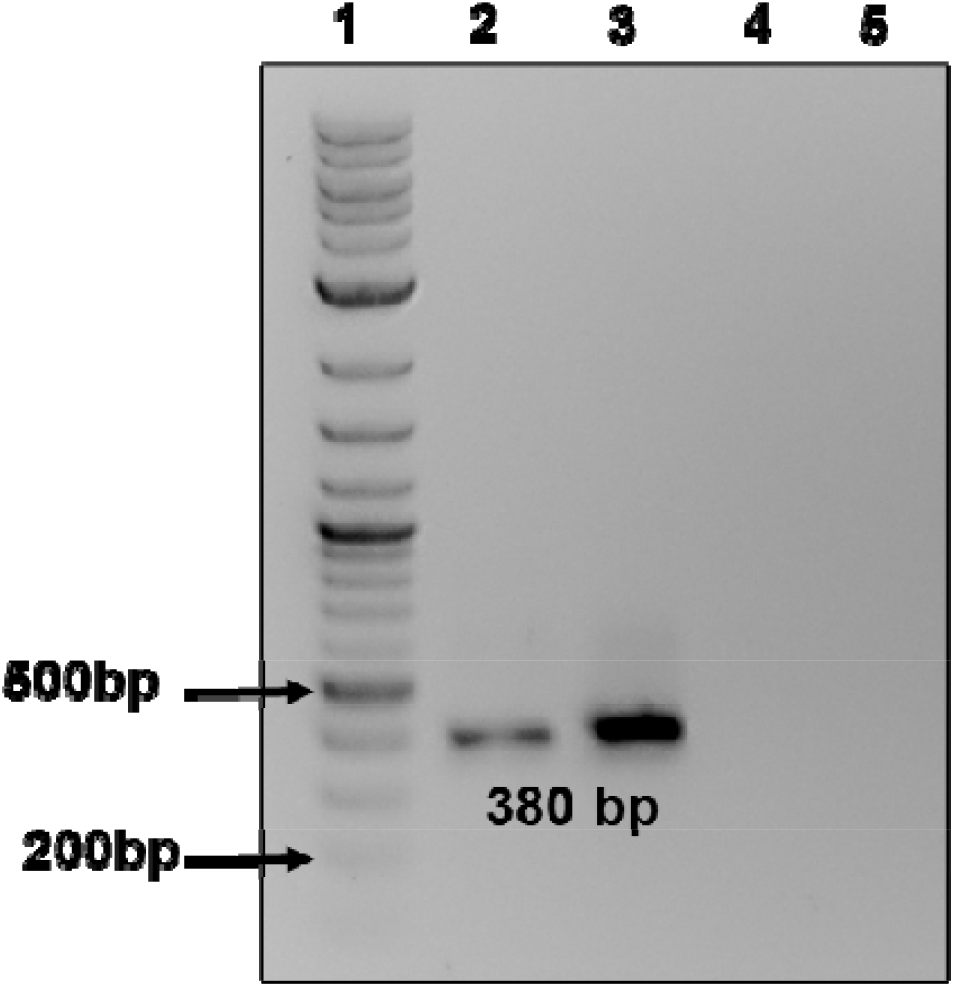
Endpoint PCR confirms Y-chromosome gene *GUY1* is present in *An. stephensi* cells. Amplicons from *An. stephensi* gDNA using *GUY1*-specific primers. Lane 1, DNA ladder; Lane 2, *An. stephensi* gDNA samples from cells; Lane 3, gDNA from *An. stephensi* male mosquito sample; Lane 4, gDNA from *An. stephensi* female mosquito sample; Lane 5, no template control (NTC).

Together, these results confirm that the cultured cells maintain the expected diploid karyotype and at least some are of male origin, as evidenced by both cytogenetic and molecular markers.

### Transfection reagent comparison in *An. stephensi* cells

To identify an efficient transfection system for *An. stephensi* cells, three commercially available reagents (Lipofectamine 3000, Lipofectamine LTX, and TransIT-PRO) were evaluated. Transfection efficiency was assessed using plasmid AGG2463, which expresses ZsGreen under the control of the 3xP3 promoter. At 48 h post-transfection, efficiency was examined by fluorescence microscopy.

Lipofectamine 3000 was tested using 2.5 μg of DNA per well with either 3.75 μL or 7.5 μL of reagent as recommended by the manufacturer for optimisation. Under both conditions, only a small number of fluorescent cells were observed, with weak and sparse signal, indicating low transfectability in *An. stephensi* cells (**Additional file 2: Figure S2**).

Similarly, Lipofectamine LTX was evaluated using 2 μg of DNA per well with 5 μL, 10 μL, or 15 μL of reagent. Across all tested conditions, the number of ZsGreen-positive cells remained minimal, suggesting poor compatibility with this cell line (**Additional file 3: Figure S3**).

In contrast, TransIT-PRO resulted in detectable transfection across all DNA concentrations tested (1.25 μg, 2.5 μg, and 5 μg per well), using a constant reagent-to-DNA ratio supplemented with TransIT-Boost. The number of fluorescent cells and overall signal intensity were highest at 2.5 μg DNA, indicating optimal transfection under these conditions (Fig 4C). Increasing the DNA amount to 5 μg did not lead to a noticeable increase in the number of transfected cells (Fig 4D), while 1.25 μg DNA produced fewer fluorescent cells with weaker signal (Fig. 4 B).

**Figure 4.**
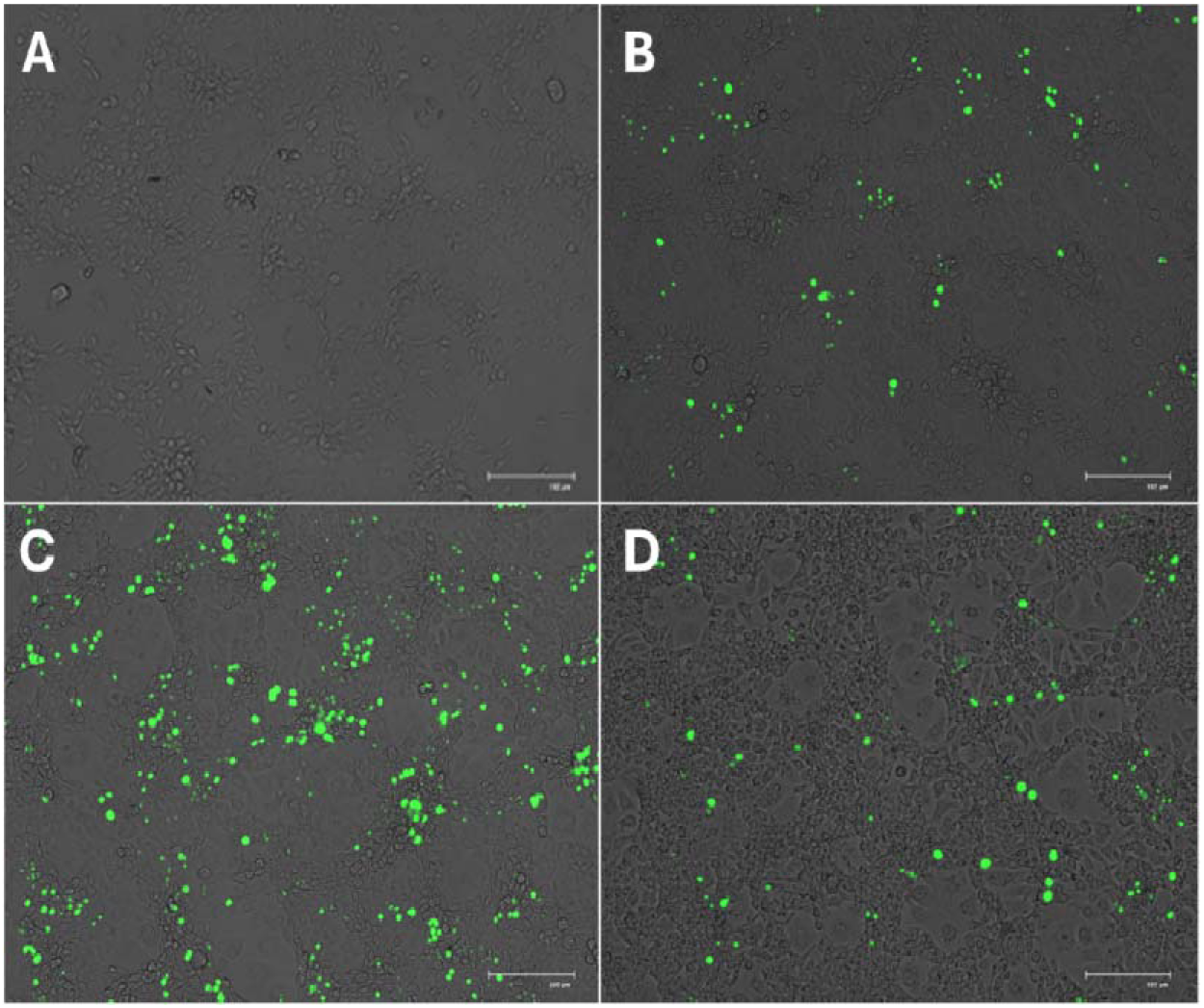
*An. stephensi* cells can be transfected using TransIT-PRO. ZsGreen fluorescence microscopy images of cells 48 h after transfection with 0 µg (A)1.25 µg (B), 2.5 µg (C), and 5 µg (D) per well.

**Figure 5.**
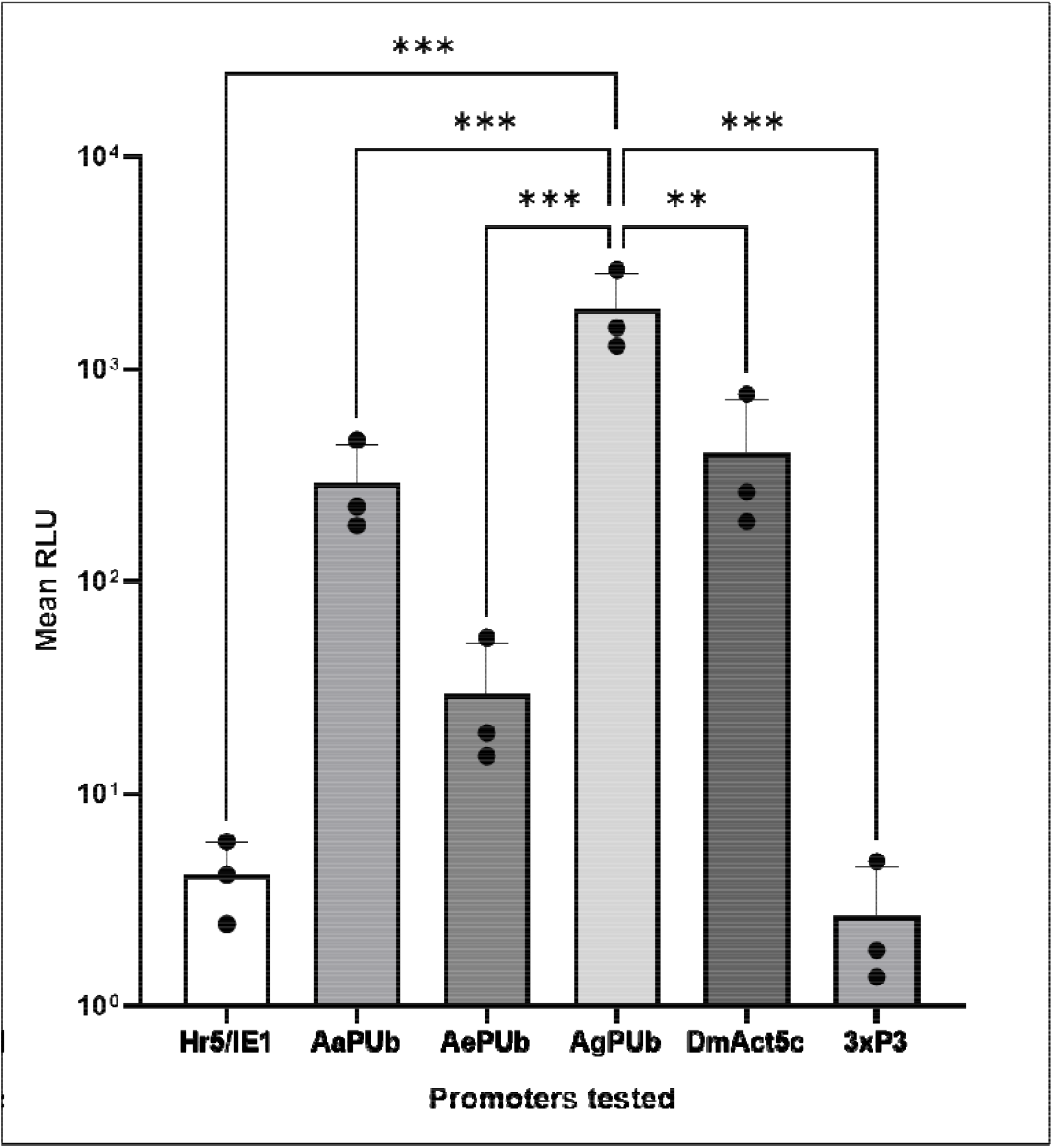
Promoter activity in *An. stephensi* cells measured by dual-luciferase assay. NanoLuc luciferase expression was driven by five Pol II promoters: Hr5/IE1, Aedes aegypti PUb (AePUb), Aedes albopictus PUb (AaPUb), Anopheles gambiae PUb (AgPUb), Drosophila melanogaster Act5C (DmAct5C) and 3xP3, and normalized against Renilla luciferase expressed from the OpIE2 constitutive promoter as an internal control. Bars represent the mean firefly luciferase normalised to Renilla relative luciferase expression unit (RLU) ± SD from three independent transfections of 8 technical replicates. Statistical significance was assessed using one-way ANOVA followed by Dunnett’s multiple comparisons test, with each promoter compared to AgPUb. Pairwise comparisons are presented in Supplementary Table S1. *p < 0.05, **p < 0.01, ***p < 0.001.

Overall, TransIT-PRO consistently yielded a higher number of ZsGreen-positive cells and stronger fluorescence compared to both Lipofectamine 3000 and Lipofectamine LTX. These findings were consistently reproduced across independent experiments, underscoring the robustness and reliability of the optimized transfection conditions in *An. stephensi* cells.

### Dual-luciferase reporter assay for Pol II promoter activity in *An. stephensi* cells

We transfected *An. stephensi* cells in 96-well plates using TransIT-PRO reagent at 2.5 μL per μg of DNA, based on prior optimization experiments. Each construct carried Firefly luciferase under the control of a test RNA polymerase II promoter: Hr5/IE1, 3xP3, *Drosophila melanogaster Act5C* (DmAct5C), *Aedes aegypti polyubiquitin* (AePUb), *Aedes albopictus polyubiquitin* (AaPUb), or *Anopheles gambiae polyubiquitin* (AgPUb). A control construct expressing Renilla luciferase driven by the OpIE2 promoter was used as an internal normalization control to compensate for variations in transfection efficiency and cell viability.

Statistical analysis of the primary dataset was performed using one-way ANOVA followed by Dunnett’s multiple comparisons test, using AgPUb as the reference promoter. AgPUb exhibited significantly higher transcriptional activity compared to Hr5/IE1, AePUb, AaPUb, DmAct5C, and 3xP3 (Fig.5; p < 0.01 for all comparisons).

To further evaluate promoter relationships, pairwise comparisons were additionally performed using one-way ANOVA followed by Tukey’s multiple comparisons test. This analysis confirmed that AgPUb was significantly higher than all other promoters tested (e.g., vs Hr5/IE1 p = 0.0006; vs AePUb p = 0.0007; vs AaPUb p = 0.0024; vs DmAct5C p = 0.0044; vs 3xP3 p = 0.0006). Significant differences were also observed among selected non-reference promoter comparisons, including AgPUb vs DmAct5C and AgPUb vs 3xP3Full statistical analysis, including one-way ANOVA and Tukey’s/Dunnett’s multiple comparisons tests, is provided in **Additional file 4: Table S1**.

Overall, AgPUb demonstrated the highest promoter activity in *An. stephensi* cells, with DmAct5C and AePUb showing moderate activity, while Hr5/IE1, AaPUb, and 3xP3 exhibited low transcriptional activity.

## Discussion

Robust *Anopheles*-derived cell lines provide a useful tool for advancing molecular and genetic studies in malaria vectors, however there is a paucity of well-characterized *Anopheles*-derived cellular models compared with those from other mosquito genera. Embryo-derived cultures have historically served as a productive source for continuous mosquito cell lines, with early work on *Aedes aegypti* embryonic explants yielding proliferative cultures exhibiting epithelial-like morphology and stable growth over serial passages (22). In general, embryo-derived cell lines are considered advantageous because they are more likely to retain native physiological and chromosomal characteristics, thereby improving their relevance as experimental systems (10). This stands in contrast to long-established immortalized mosquito cell lines, which may gradually accumulate genomic alterations and lose key chromosomal features over time (11,23). Nevertheless, the utility of a newly established cell line extends beyond its stability; its value ultimately depends on its ability to support transgene expression and genome manipulation. In this context, evaluating regulatory elements represents a crucial next step in its functional characterisation.

Efficient transfection is essential for functional characterization of new cell lines and genetic manipulation, so we compared three commercial reagents to identify an optimal system for *An. stephensi* cells. Lipofectamine 3000 and Lipofectamine LTX showed very low efficiency, with only rare ZsGreen-positive cells despite exploring different DNA and reagent concentrations, consistent with previous reports that lipid-based reagents optimized for mammalian cells often perform poorly in insect systems. As noted by Kost & Condreay (24), differences in membrane composition and endocytic pathways between insect and mammalian cells can strongly limit uptake and expression, and similar limitations have been described in other mosquito-derived lines where standard reagents require substantial optimization (9,25). In contrast, TransIT-PRO showed markedly improved performance, with consistent reporter expression across conditions and an optimum at 2.5 μg DNA, after which efficiency plateaued, in line with general observations that transfection systems often reach a saturation point beyond which additional DNA does not improve uptake or expression. Overall, TransIT-PRO’s broader formulation appears better suited for more refractory insect cells, reinforcing the need for empirical optimization in non-mammalian systems. The ability to achieve reliable transgene expression is particularly important for downstream applications such as promoter analysis, gene function studies, and genome editing approaches, including CRISPR/Cas-based systems (26,27). In this context, the identification of TransIT-PRO as an effective reagent represents a key step in enabling functional genomic studies in this newly established *An. stephensi* cell line.

In this study, we evaluated the activity of several RNA polymerase II promoters in *An. stephensi* cells using a dual-luciferase reporter assay. Our results demonstrate that promoter choice has a substantial impact on transgene expression in this system. The *Anopheles gambiae polyubiquitin promoter* (AgPUb) exhibits stronger activity compared to all other promoters tested, consistent with previous reports highlighting the robustness of ubiquitin promoters in mosquitoes, particularly those derived from closely related species (13,28). The *Drosophila melanogaster Act5C* promoter showed moderate activity in *An. stephensi* cells, indicating that heterologous insect promoters can function across species boundaries, albeit with reduced efficiency. As a widely used constitutive promoter driving actin expression in insect systems (28,29), its performance here confirms it as a reliable option for moderate transgene expression in mosquito cells. The *Aedes aegypti polyubiquitin* promoter exhibited intermediate activity, although its performance was not significantly different from lower-expressing constructs, suggesting context-dependent variability. This may reflect partial compatibility of ubiquitin-driven transcriptional regulation in *An. stephensi*, where promoter strength appears less robust than in its native species. The *Aedes albopictus polyubiquitin* promoter, 3xP3, and Hr5/IE1 promoters all showed consistently low activity in this system. The weak performance of the AePUb promoter likely reflects species-specific regulatory divergence. The 3xP3 promoter is expected to be limited in cultured cells due to its tissue-specific design (15). While the Hr5/IE1 promoter has previously been reported to drive robust transgene expression in several insect and mosquito systems, including in the context of synthetic biology applications, its activity in the present study was comparatively low, suggesting that promoter performance may be strongly influenced by cellular context and species-specific transcriptional machinery (30–33). Overall, these results highlight that promoter performance in *An. stephensi* is highly context dependent, reinforcing the importance of empirical validation for mosquito molecular tools.

Cytogenetic analysis of the established *An. stephensi* cell line provides critical insight into its chromosomal stability and genetic composition. The observed diploid complement of 2n = 6 aligns with classical karyotypic descriptions of *An. stephensi* (20,34), indicating that chromosomal integrity has been maintained during the early establishment and propagation of the cell line. Among the chromosomes, one smaller and morphologically distinct element suggested the presence of a Y chromosome. This finding is noteworthy, as the retention of sex chromosomes, particularly the Y, can be compromised in long-term or immortalized insect cell lines (35,36).To confirm its identity, PCR amplification targeting the Y-linked *AsGUY1* gene was performed. *AsGUY1* is a well-characterized male-specific marker in *An. stephensi* exclusively localized to the Y chromosome (18). Successful amplification of this gene confirmed not only the presence of the Y chromosome but also the male origin of at least a subset of the cultured cells. This is particularly relevant for research into sex determination, male-specific gene function, and the development of vector control strategies, including sex ratio distortion and male-targeted gene drives (37).

In conclusion, this study presents the development and initial characterization of a novel *An. stephensi* cell line and demonstrates its suitability for initial functional genomic assays. Through systematic evaluation, we confirmed that the cell line is amenable to transfection using multiple commercially available reagents and is compatible with RNA polymerase II– driven gene expression. Despite over two decades of progress in mosquito transgenesis, the repertoire of well-characterized promoter elements remains limited (25,38). Addressing this gap, we assessed the activity of several Pol II promoters—including the synthetic 3xP3 and polyubiquitin promoters from *Anopheles gambiae, Aedes aegypti*, and *Anopheles albopictus*—and demonstrated their functionality using a dual-luciferase reporter system. In addition, karyotypic analysis confirmed chromosomal integrity and supported accurate species identification, underscoring the stability of the cell line. Collectively, these findings indicate that this *An. stephensi* cell line can be transfected and supports RNA polymerase II-driven gene expression.

## Materials and Methods

### Establishment of the cell line

*Anopheles stephensi* SDA-500 strain embryos were collected and aged 24 hours. The embryos were then collected into a cell strainer and surface decontaminated by submerging in 10% sodium hypochlorite solution for 5 minutes, then rinsing twice with sterile water. They were then submerged in 70% ethanol for 2.5 minutes, followed by two water washes. Embryos were then collected into a 1.5 ml tube with a paintbrush and homogenised in 500 µl of complete media (Leibovitz’s L-15 media supplemented with 20% foetal bovine serum (FBS), 10% tryptose phosphate broth, 1% penicillin–streptomycin, 1% gentamycin) using a sterile pestle. The homogenate was passed through a new cell strainer (Corning 100 µm). Cells were plated into T-25 flasks and maintained at 28□°C in a humid environment without added CO_2_. Each week, 50% of the media was replaced. After one month the media was completely replaced weekly. After approximately 6 months the flask became confluent and was split 1:2. The cells continued to be split 1:2 when confluent, and the frequency increased over time until they stabilised being passed ∼1:5 twice weekly. At this point (passage 15, approximately one year after initial culture) the FBS was reduced to 10% and the gentamycin removed from the media. The culture was maintained for more than 40 passages.

### COI barcoding PCR

Genomic DNA was isolated from *An. stephensi* cell cultures using the Nucleospin Tissue DNA extraction kit (Machery-Nagel), following the manufacturer’s protocol. A ∼658 bp fragment of the mitochondrial cytochrome c oxidase subunit I (COI) gene was amplified by polymerase chain reaction (PCR) using the universal barcoding primers LCO1490 (5′-GGTCAACAAATCATAAAGATATTGG-3′) and HCO2198 (5′-TAAACTTCAGGGTGACCAAAAAATCA-3′) (19). PCR reactions were carried out in a total volume of 25 µL using 1× DreamTaq PCR Master Mix (2X) (Thermo Fisher Scientific), prepared according to the manufacturer’s instructions. Each reaction included approximately 50 ng of genomic DNA as the template. The thermal cycling profile consisted of an initial denaturation at 94°C for 3 minutes; 35 cycles of 94°C for 30 seconds, 50°C for 30 seconds, and 72°C for 1 minute; followed by a final extension at 72°C for 10 minutes. PCR products were purified using the Nucleospin Gel and PCR clean-up kit (Machery-Nagel) and submitted for Sanger sequencing using the same primers. Sequence chromatograms were assembled and used for species identification via BLASTn analysis against the NCBI nucleotide database.

### Transfections

*An. stephensi* cells were harvested and resuspended in L15-complete medium at a density of 4 × 10□ cells/mL, as determined using a Scepter 2.0 automated cell counter (Millipore). A total of 2 mL of the cell suspension (equivalent to 8 × 10□ cells) was seeded into each well of a 6-well plate and incubated at 28 °C under standard culture conditions for 24 h to allow cell adherence and growth, reaching 90-95% confluence before transfection. To evaluate transfection efficiency, three commercial transfection reagents: Lipofectamine 3000, Lipofectamine LTX, and TransIT-Pro, were tested at varying reagent concentrations. Transfections were performed in six well plates using 2.5 µg of AGG2463 plasmid DNA per well. The AGG2463 plasmid expresses the fluorescent reporter gene ZsGreen under the control of the 3xP3 promoter. DNA-reagent complexes were prepared according to the manufacturer’s protocols in Opti-MEM I medium. Cell transfection was assessed 48 hours post-transfection by fluorescence microscopy using a Floid Cell Imaging Station (Life Technologies) (20x objective).

### Lipofectamine 3000 (Thermo Fisher Scientific)

For each well, 2.5 µg of AGG2463 plasmid DNA was mixed with 10 µL of P3000 reagent in Opti-MEM to a final volume of 250 µL. In parallel, Lipofectamine 3000 was diluted in 125 µL of Opti-MEM at two different amounts (either 3.75 µL or 7.5 µL) to test two reagent concentrations. The diluted DNA and Lipofectamine 3000 solutions were combined, incubated at room temperature for 10–15 min, and then added to cells maintained in L-15 complete medium.

### TransIT-Pro (Mirus Bio)

Transfections were performed using varying amounts of plasmid DNA (1.25, 2.5, or 4.9 µg per well). Plasmid DNA was combined with 2.5 µL of Boost reagent and 5 µL of TransIT-Pro in Opti-MEM to a final volume of 250 µL. The transfection mixture was incubated at room temperature for 20 min and then added to the cells maintained in L-15 complete medium.

### Lipofectamine LTX (Thermo Fisher Scientific)

For transfections, 2 µg of plasmid DNA was mixed with 2.5 µL of PLUS reagent in Opti-MEM to a final volume of 250 µL. In parallel, Lipofectamine LTX was diluted in 150 µL of Opti-MEM at three different volumes (5, 10, or 15 µL) to test varying reagent concentrations. The diluted DNA and Lipofectamine LTX solutions were combined, incubated at room temperature for 5–10 min, and then added to cells maintained in L-15 complete medium.

Following transfection, cells were incubated for an additional 48 hours at 28□°C and approximately 75□% relative humidity. Transfection efficiency was assessed by fluorescence microscopy through evaluation of ZsGreen-derived green fluorescence, including both signal intensity and distribution within the cells. Negative controls consisted of untransfected cells and reagent-only conditions, and all experimental conditions were performed in triplicate.

### Dual-luciferase assay

*An. stephensi* cells were seeded into 96-well plates at a density of 5 x 10□ cells per well and incubated for 24 h. Cells were then co-transfected with two plasmids: (i) a test construct encoding firefly luciferase driven by a single RNA polymerase II (Pol II) promoter— Hr5/IE1 (AGG1185), *Aedes aegypti polyubiquitin* (AePUb; AGG1747), *Aedes albopictus polyubiquitin* (AaPUb: AGG1264), 3xP3 (AGG2894), *Drosophila melanogaster Act5C* (DmAct5C; AGG2895), or *Anopheles gambiae polyubiquitin* (AgPUb; AGG2896)—at 80 ng per well, and (ii) an internal control plasmid expressing Renilla luciferase under the control of the OpIE2 promoter with an SV40 polyadenylation signal (AGG1080; 10 ng per well). Plasmids AGG1185, AGG1747 (OR236189), and AGG1264 were obtained from existing laboratory stocks; AGG2894, AGG2895, and AGG2896 were synthesized commercially by Genscript.

Transfections were performed using TransIT-PRO Transfection Reagent (Mirus Bio) under conditions optimized for *An. stephensi* cells. Following transfection, cells were incubated at 28 °C for 48 h. Cells were lysed in passive lysis buffer and Firefly and Renilla luciferase activities were measured using the Dual-Glo Luciferase Assay System (Promega), and luminescence was quantified with a GloMax Multi-Detection System (Promega). Each condition was analysed using eight technical replicates and three independent biological replicates.

### Chromosome Preparation and Karyotyping

*An. stephensi* cells were cultured in L-15 medium (as described above) at 28 °C until reaching 70–80% confluency. To arrest cells in metaphase, cultures were treated with colcemid (0.1 µg/mL) for 2 h. Cells were harvested by centrifugation at 1000 × g for 5 min, resuspended in 0.075 M KCl for hypotonic treatment (20 min, room temperature), and fixed in freshly prepared Carnoy’s fixative (3:1 methanol: acetic acid). The fixation step was repeated three times to ensure optimal preservation. Droplets of fixed cells were air-dried on clean glass slides. Slides were stained with 10% Giemsa for 15 min, rinsed with distilled water, and mounted with DPX (Distrene, Plasticizer, and Xylene). Chromosomes were examined using an EVOS M7000 imaging system at 40x magnification, and representative images were captured for analysis.

### Y chromosome confirmation by PCR using AsGUY1 primers

Genomic DNA was extracted from *An. stephensi* cells using the NucleoSpin Tissue Kit (Macherey-Nagel) according to the manufacturer’s instructions. PCR amplification was performed using primers specific to AsGUY1 (F: 5′-GGGCATCAAGCCTCCCACCA-3′ and R: 5′-TGTTGTGTGCGTGTCCGACCTA-3), a male-specific gene located on the Y chromosome of *An. stephensi* (18). PCR reactions were prepared in a final volume of 25 µL containing 12.5 µL Q5 High-Fidelity 2X Master Mix, 0.5 µL of each primer (10 µM), 1–2 µL genomic DNA template, and nuclease-free water to volume. Thermal cycling was carried out using the following program: initial denaturation at 98 °C for 30 s, followed by 35 cycles of denaturation at 98 °C for 10 s, annealing at 60 °C for 20 s, and extension at 72 °C for 30 s, with a final extension at 72 °C for 2 min. Amplified products were separated by agarose gel electrophoresis and visualized using a nucleic acid staining method.

### Statistics

Statistical analysis was performed in Graphpad Prism 9.4.1 using one-way ANOVA to test for differences in promoter activity among groups, followed by Tukey’s post hoc test for pairwise comparisons, with significance set at p < 0.05.

## Acknowledgements

This work was funded, in whole or in part, by the Gates Foundation [INV-008549]. The conclusions and opinions expressed in this work are those of the author(s) alone and shall not be attributed to the foundation. Under the grant conditions of the foundation, a Creative Commons Attribution 4.0 License has already been assigned to the Author Accepted Manuscript version that might arise from this submission. Please note works submitted as a preprint have not undergone a peer review process.

## Authors’ contributions

L.A., M.A., and SK. developed the research plan and experimental strategy. M.A., S.K., and D.J. performed the experiments. S.K. analyzed the data. S.K. and M.A. drafted the manuscript; M.A. and L.A. revised the manuscript. All authors read and approved the manuscript.

## Declarations

## Ethics approval and consent to participate

Not applicable.

## Consent for publication

Not applicable.

## Additional file 1 Figure S1

Amplification and sequencing of the mitochondrial COI gene from SDA-500 cells

Amplified Anopheles stephensi gDNA sample: Lane 1, DNA ladder; Lane 2, no template control; Lane 3

## Additional file 2 Figure S2

An. stephensi cells can be transfected using Lipofectamine 3000.

ZsGreen fluorescence microscopy images of cells 48 h after transfection with 0 µL (A), 3.75 µL (B) and 7.5 µL (C) per well.

## Additional file 3 Figure S3

An. stephensi cells can be transfected using Lipofectamine LTX

ZsGreen fluorescence microscopy images of cells 48 h after transfection with 0 µL (A), 5 µL (B), 10 µL (C) and 15 µL (D) of reagent per well.

## Additional file 4 Table S1

Statistical comparisons of promoter transcriptional activity

One-way ANOVA and Tukey’s/Dunnett’s multiple comparisons tests. Adjusted p-values were used to determine statistical significance. ns, not significant; *, p < 0.05; **, p < 0.01; ***, p < 0.001.

