## Additional file 1: Figure S1 for "Development and validation of an SDA-500 *Anopheles stephensi* cell line for molecular studies"

Figure S1. Amplification and sequencing of the mitochondrial COI gene from SDA-500 cells.

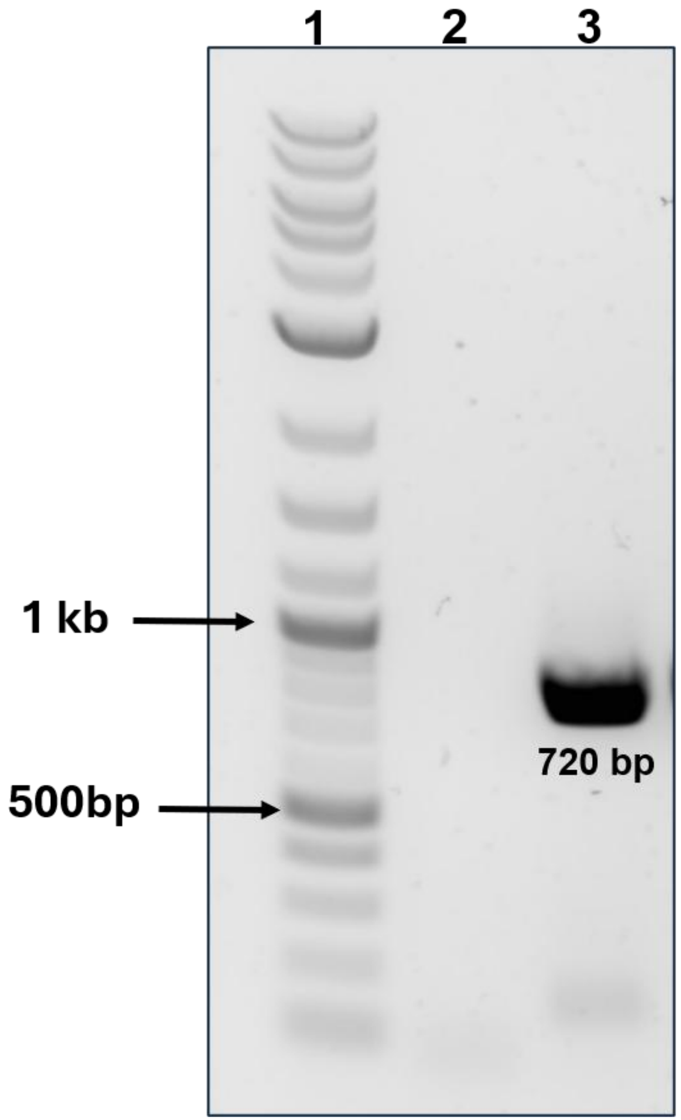


Lane 1, DNA ladder; Lane 2, no template control; Lane 3, amplified *Anopheles stephensi* gDNA samples
