## Additional file 2 Figure S2 for "Development and validation of an SDA-500 *Anopheles stephensi* cell line for molecular studies"

Figure S2. *An. stephensi* cells can be transfected using Lipofectamine 3000

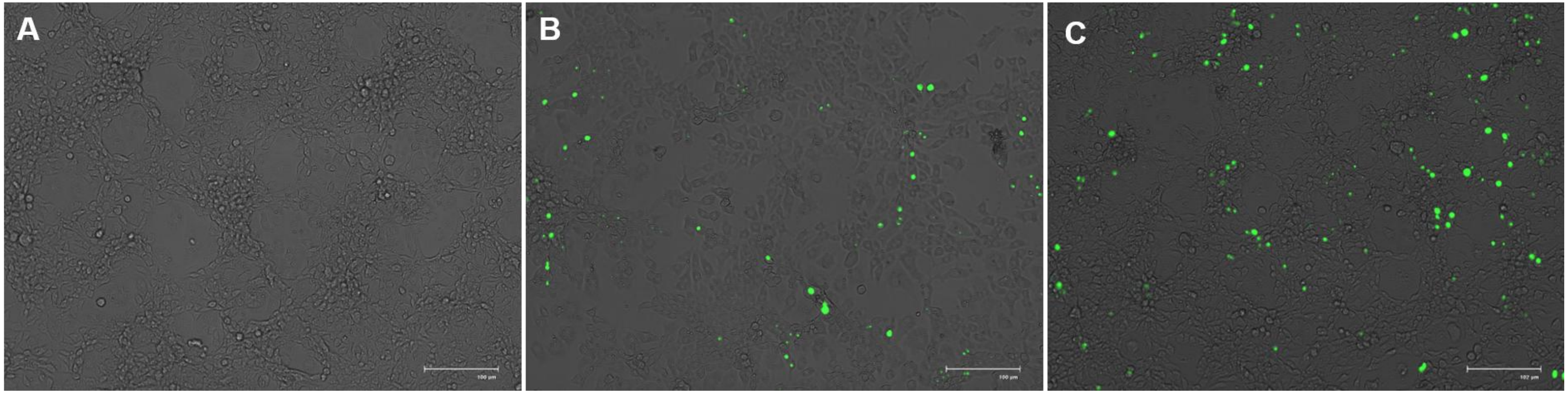


ZsGreen fluorescence microscopy images of cells 48 h after transfection with 0 µL (A), 3.75 µL (B) and 7.5 µL (C) per well.
