## Additional file 3 Figure S3 for "Development and validation of an SDA-500 *Anopheles stephensi* cell line for molecular studies"

### *An. stephensi* cells can be transfected using Lipofectamine LTX

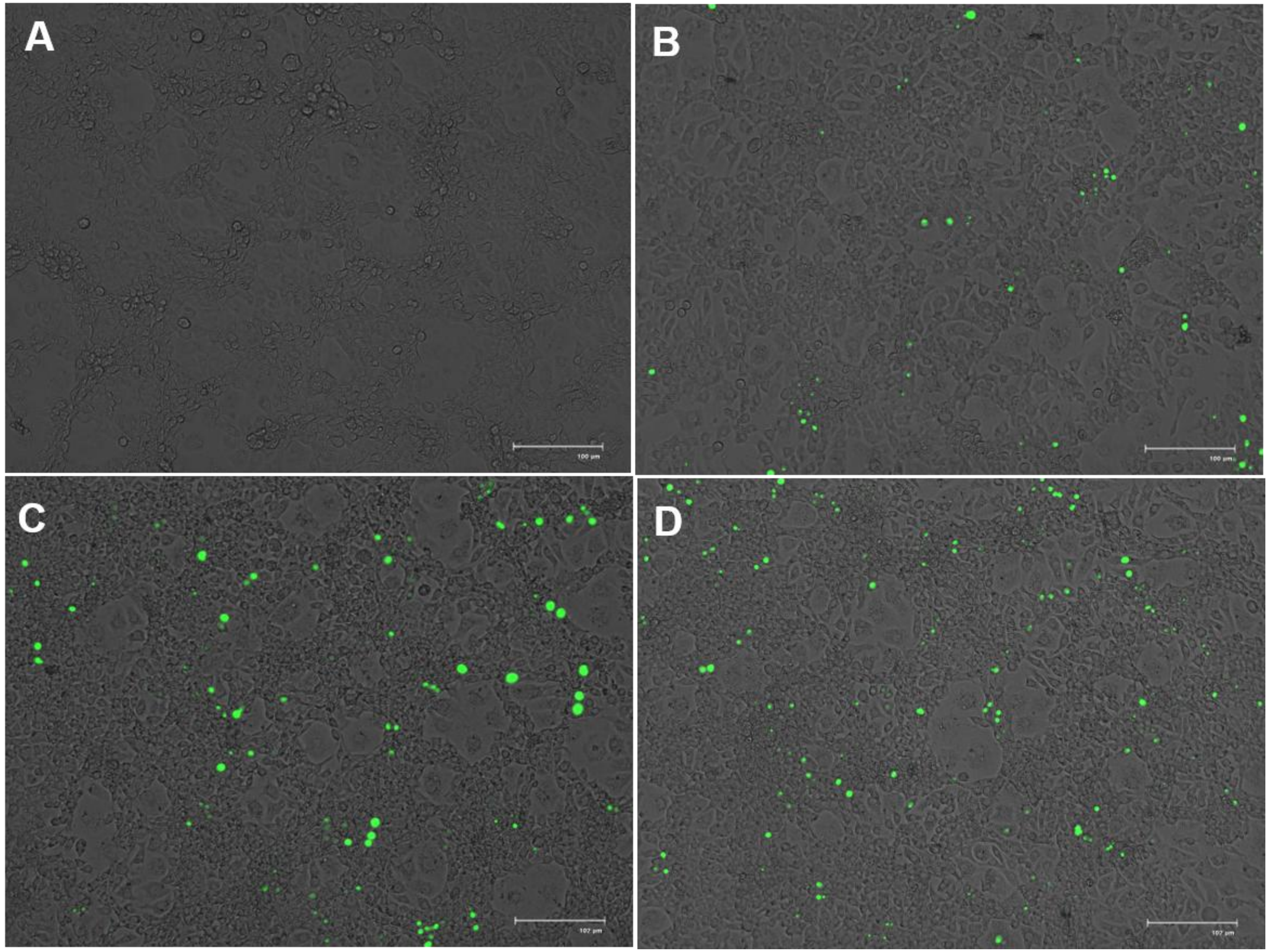


ZsGreen fluorescence microscopy images of cells 48 h after transfection with 0 µL (A), 5 µL (B), 10 µL (C) and 15 µL (D) of reagent per well.
