## Additional file 4 Table S1 for "Development and validation of an SDA-500 *Anopheles stephensi* cell line for molecular studies"

**Statistical comparisons of promoter transcriptional activity**

| **Comparison** | **Test** | **Mean difference** | **Adjusted p-value** | **Significance** |
| --- | --- | --- | --- | --- |
| HR5-IE1 vs AaPUb | Tukey | -287.5 | 0.9376 | ns |
| HR5-IE1 vs AePUb | Tukey | -25.56 | >0.9999 | ns |
| HR5-IE1 vs AgPUb | Tukey | -1931 | 0.0006 | *** |
| HR5-IE1 vs DmAct5C | Tukey | -403.2 | 0.7947 | ns |
| HR5-IE1 vs 3xP3 | Tukey | 1.517 | >0.9999 | ns |
| AaPUb vs AePUb | Tukey | 261.9 | 0.9569 | ns |
| AaPUb vs AgPUb | Tukey | -1644 | 0.0024 | ** |
| Aa pUb vs. Dm Act 5c | Tukey | -115.7 | 0.9989 | ns |
| Aa pUb vs. 3X P3 | Tukey | 289.0 | 0.9363 | ns |
| Ae pUb vs. Ag pUb | Tukey | -1906 | 0.0007 | *** |
| Ae pUb vs. Dm Act 5c | Tukey | -377.6 | 0.8334 | ns |
| Ae pUb vs. 3X P3 | Tukey | 27.08 | >0.9999 | ns |
| Ag pUb vs. Dm Act 5c | Tukey | 1528 | 0.0044 | ** |
| Ag pUb vs. 3X P3 | Tukey | 1933 | 0.0006 | *** |
| Dm Act 5c vs. 3X P3 | Tukey | 404.7 | 0.7923 | ns |
| Ag pUb vs. HR5-IE1 | Dunnett | 1931 | 0.0003 | *** |
| Ag pUb vs. Aa pUb | Dunnett | 1644 | 0.0010 | *** |
| Ag pUb vs. Ae pUb | Dunnett | 1906 | 0.0003 | *** |
| Ag pUb vs. Dm Act 5c | Dunnett | 1528 | 0.0018 | ** |
| Ag pUb vs. 3X P3 | Dunnett | 1933 | 0.0003 | *** |

one-way ANOVA and Tukey’s/Dunnett’s multiple comparisons tests. Adjusted p-values were used to determine statistical significance. ns, not significant; *, p < 0.05; **, p < 0.01; ***, p < 0.001.
